# MS2-TRIBE evaluates protein-RNA interactions and nuclear organization of transcription by RNA editing

**DOI:** 10.1101/829606

**Authors:** Jeetayu Biswas, Reazur Rahman, Varun Gupta, Michael Rosbash, Robert H. Singer

**Affiliations:** Department of Anatomy and Structural Biology, Albert Einstein College of Medicine, Bronx, NY, 10461, USA; Department of Biology, Howard Hughes Medical Institute and National Center for Behavioral Genomics, Brandeis University, Waltham, MA, 02453, USA; Department of Cell Biology, Albert Einstein College of Medicine, Bronx, NY, 10461, USA; Howard Hughes Medical Institute, Janelia Research Campus, Ashburn, VA, 20147, USA

**Keywords:** CLIP, TRIBE, Adenosine Deaminase, RNA, Next Generation Sequencing, RNA Binding Proteins, transcription domains

## Abstract

Nearly every step of RNA regulation is mediated by binding proteins (RBPs). The most common method to identify specific RBP target transcripts in vivo is by crosslinking (“CLIP” and its variants), which rely on protein-RNA crosslinking and specific antibodies. Another recently introduced method exploits RNA editing, with the catalytic domain of ADAR covalently attached to a specific RBP (“TRIBE”). Both approaches suffer from difficulties in distinguishing real RNA targets from false negative and especially false positive signals. To critically evaluate this problem, we used fibroblasts from a mouse where every endogenous β-actin mRNA molecule was tagged with the bacteriophage MS2 RNA stem loops; hence there is only a single bona fide target mRNA for the MS2 capsid protein (MCP). CLIP and TRIBE could both detect the single RNA target, albeit with some false positives (transcripts lacking the MS2 stem loops). Consistent false positive CLIP signals could be attributed to nonspecific antibody crosslinking. To our surprise, the supposed false positive TRIBE targets correlated with the location of genes spatially proximal to the β-actin gene. This result indicates that MCP-ADAR bound to β-actin mRNA contacted and edited nearby nascent transcripts, as evidenced by frequent intronic editing. Importantly, nascent transcripts on nearby chromosomes were also edited, agreeing with the interchromosomal contacts observed in chromosome paint and Hi-C. The identification of nascent RNA-RNA contacts imply that RNA-regulatory proteins such as splicing factors can associate with multiple nascent transcripts and thereby form domains of post-transcriptional activity, which increase their local concentrations. These results more generally indicate that TRIBE combined with the MS2 system, MS2-TRIBE, is a new tool to study nuclear RNA organization and regulation.

## Introduction

RNAs are regulated throughout all aspects of their lives (Vera et al., 2016). Their regulation by RNA binding proteins (RBPs) provides spatiotemporal control over processing, export, localization, translation and decay. RBP genes make up approximately 10% of all protein coding genes in the human genome (Gerstberger et al., 2014), and RBP families have coevolved with their RNA targets to provide control over functionally related transcripts (Hogan et al., 2015). Many RBPs contain more than one RNA binding domain, with each domain contributing to the final specificity of the protein (Gerstberger et al., 2014). Accurately defining the targets of a given RBP has been a longstanding challenge in biology due to their complex and multivalent interactions.

*In vitro* approaches have determined the sequence specificity of single domains within RBPs. Approaches such as SELEX and “RNA bind-n-seq” (Lambert et al., 2014) have determined the consensus sequence preference for RBPs using short RNA fragments (<40nt). Complementary approaches have found consensus binding sites *in vivo*, using crosslinking and immunoprecipitation (CLIP).

Cell lysis prior to immunoprecipitation promotes adventitious interactions such as those between cytoplasmic RBPs and nuclear RNAs (Mili and Steitz, 2004). To overcome these limitations, CLIP seq and its derivatives use UV to irreversibly crosslink protein and RNA within cells (Ule et al., 2003). These interactions can be defined with single nucleotide resolution (Zhang and Darnell, 2011). However, CLIP relies on antibodies and suffers from high background due to nonspecific antibody-antigen interactions (Friedersdorf and Keene, 2014). Therefore, an orthogonal approach is important to obtain high confidence targets for RNA binding proteins of interest. It should be highly specific, antibody independent, retain information about the total RNA present for a given species and be limited in the amount of sample and its handling steps.

Two approaches have recently been developed, RNA editing in *Drosophila* (“TRIBE”, McMahon et al., 2016) and RNA tagging in yeast (Lapointe et al., 2015). Both approaches utilize enzymes fused to an RNA binding protein of interest to deposit marks on RNA targets. We chose to focus on RNA editing because of its straightforward approach utilizing standard RNA sequencing library preparation.

Conspicuously missing from previous transcriptome-wide studies of RNA binding proteins is a positive control, which could resolve rigorously the extent of false positives: the best case scenario would be an RNA binding protein that recognizes only a single transcript within the cell. Thus, any targets identified outside of the single transcript would be defined as false positives. The specificity of the MCP MBS system provides such a tool: its capsid protein (MCP) should only bind the MS2 stem loops (MBS) inserted into a gene of interest. We used cells derived from a mouse that has the MBS inserted into the endogenous β-actin gene and used the MCP as a means to evaluate various approaches to identifying targets of RNA binding proteins (Lionnet et al., 2011a).

To validate this approach, both CLIP and TRIBE were performed on cells containing the MCP MBS system. The single β-actin mRNA labeled with the MCP-MBS system allowed for unambiguous determination of background or false positives for each technique. While nonspecific antibody interactions contributed significantly to MCP-CLIP signals, there were fewer non β-actin target mRNAs with MCP-TRIBE. Moreover, these reproducible, non-random targets correlated with their proximity to the β-actin locus on the chromosome rather than manifesting some association with β-actin mRNA. This result led to the unexpected conclusion that MCP-ADAR had edited nearby transcribing RNAs. The tethered editing enzyme can therefore define a nuclear transcriptional domain with implications for how the transcriptome may be organized in the nucleus.

## Results

### MCP-ADAR edits targets consistent with binding the β-actin-MBS

We evaluated both CLIP and TRIBE with an RBP that should recognize a single target within the transcriptome. We used mouse fibroblasts containing 24xMBS integrated into the endogenous β-actin locus (Fig. 1A, (Lionnet et al., 2011b)). mRNA was imaged with smFISH (Sup Fig. 1A), and live cells imaged with MCP-GFP (Sup Fig. 1B). Both approaches demonstrated a sufficient signal to noise ratio for single molecule detection. The sub-nanomolar kD of MCP for the β-actin-MBS in vitro (Sup Fig. 1C,D) was consistent with the high specificity of MCP for its stem loop target *in vivo*.

**Fig. 1.**
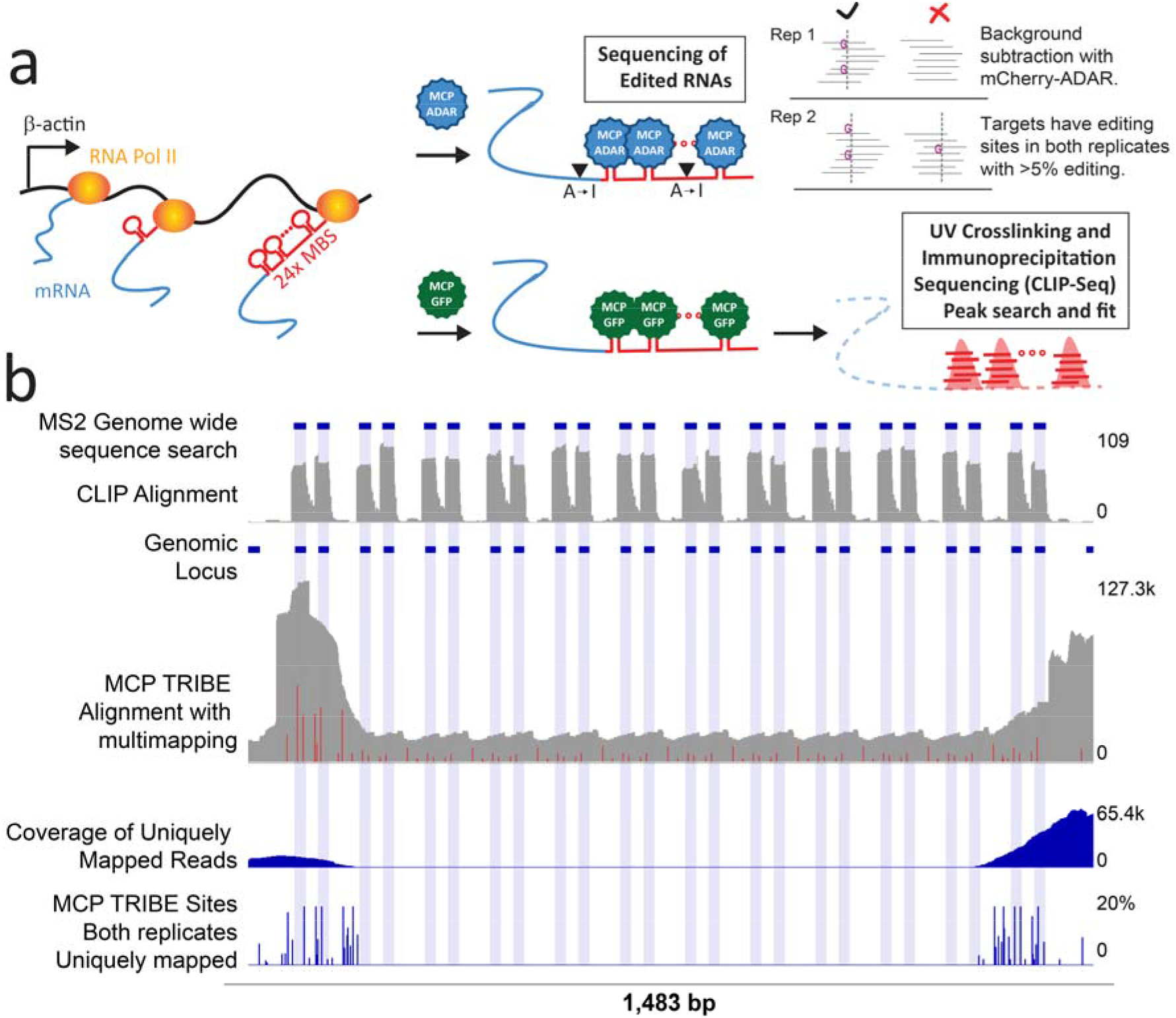
Transcriptome wide studies show CLIP peaks and TRIBE editing at β-actin-MBS. (A) Schematic of MCP wtADAR binding to MS2 stem loops that have been integrated into the endogenous β-actin locus. (Right arrows, top) Schematic of MCP ADAR binding to MS2 stem loops and editing adjacent nucleotides from adenosine to inosine (which is then recognized as guanosine upon cDNA conversion). (Right arrows, bottom) Schematic of MCP GFP binding to MS2 stem loops. UV crosslinking and immunoprecipitation (CLIP) isolates RNA fragments (red horizontal bars) which are then computationally analyzed and fit to peaks (red peaks). (B) β-actin gene, focusing on the MBS array showing (from top to bottom) MS2 sequence search, CLIP signal, genomic locus annotation, mRNA coverage with multimapping to show editing sites (red bars) within stem loop region, mRNA coverage without multimapping and editing tracks for mouse embryonic fibroblasts. Editing events are indicated by dark blue bars, average of editing percentage from both replicates, the height of the bar indicates the percentage editing at that site. Light blue shading indicate location of the stem loop nucleotides.

To determine if the tight binding of MCP to the β-actin-MBS array would lead to RNA editing of the β-actin transcript, MCP fused to ADAR was introduced into cells containing the β-actin-MBS mRNA. We also performed CLIP on cells stably expressing MCP-GFP and β-actin-MBS. Additionally, the MCP consensus binding sequence was utilized to perform an *in-silico* search for MS2 binding sites across the transcriptome (Tutucci et al., 2018). All three approaches were successful in predicting MCP binding to the target sites (Fig. 1B).

### Reproducible CLIP and TRIBE binding to β-actin-MBS

In both approaches, the β-actin-MBS array was the top ranked site (by total peak height in CLIP or by total number of editing sites in TRIBE). For example, it had a TRIBE signal 4x greater than the next most highly edited transcript in the transcriptome. This robust binding was expected since the β-actin-MBS array has the highest density of MCP binding motifs of any transcript in the transcriptome.

There were also 139 other transcripts identified with TRIBE and 105 with CLIP. One possibility is that they contained stem loop structure resembling MS2 and were due to direct binding of MCP. To address this possibility, transcripts were examined for the presence of the MCP consensus sequence and the RNAs detected by MCP TRIBE were compared to those crosslinked by MCP CLIP.

Few MCP TRIBE transcripts had showed a MCP consensus sequence, and only three transcripts were shared between MCP CLIP and MCP TRIBE in addition to β-actin (Fig. 2A). Closer inspection of the three other transcripts showed no resemblance to either the sequence or structure of MBS, suggesting that the identified transcripts are unique to each technique.

**Fig. 2.**
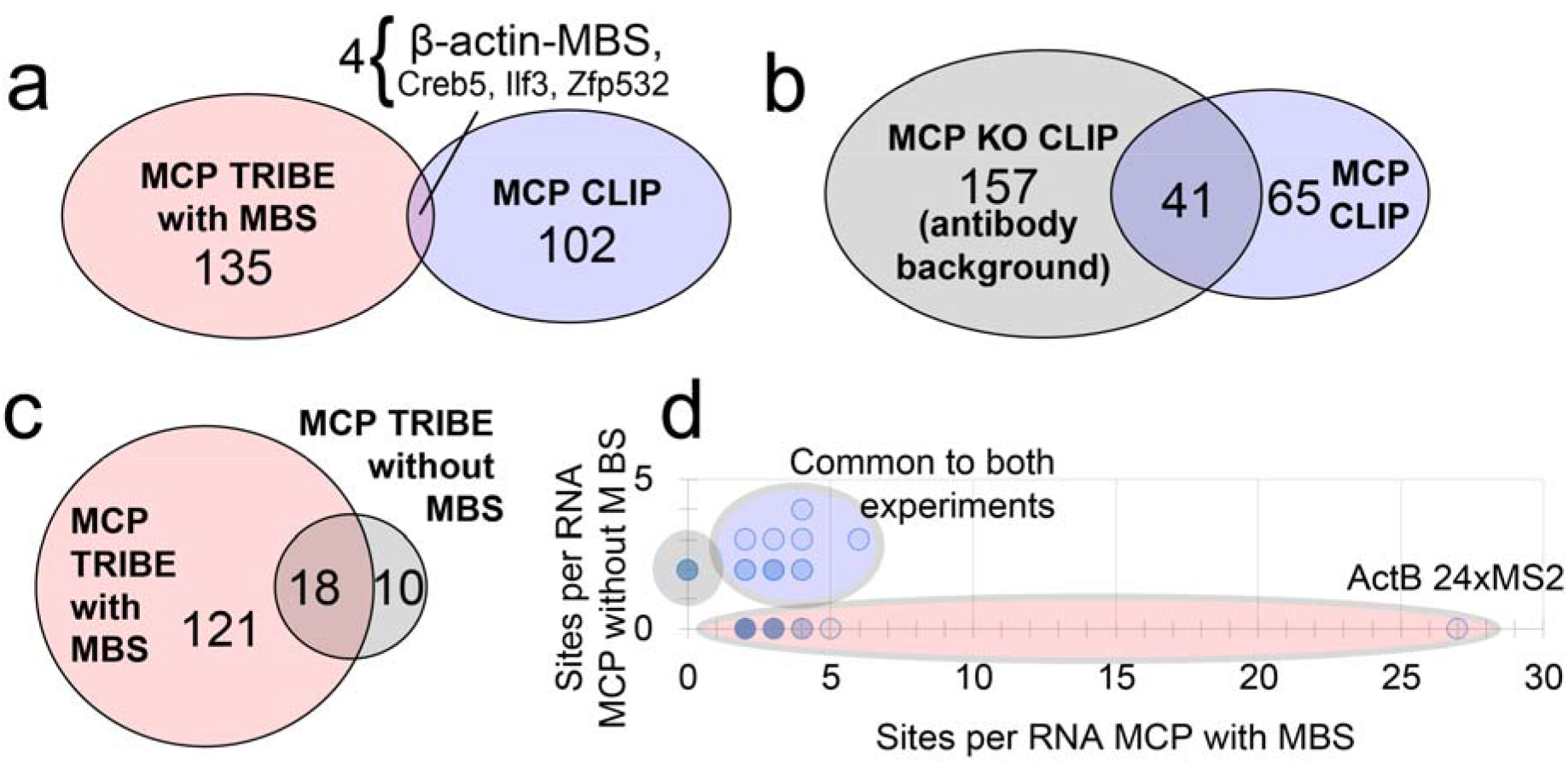
Comparison of MCP CLIP and MCP TRIBE targets. (A) Venn diagram showing significant peaks discovered in MCP CLIP (blue oval) and MCP TRIBE (red oval) recognize different target pools. Four RNAs overlap between both techniques (intersection with line to names of RNAs), with only one RNA containing evidence of the MS2 consensus sequence (β-actin-MBS). (B) Venn diagram showing intersection of RNAs found by MCP-GFP CLIP (red oval) and KO CLIP (background CLIP, blue oval). 41/106 significant CLIP peaks are reproduced in the MCP KO cells, representing antibody background. (C) Venn diagram showing Intersection of RNAs found with MCP MBS TRIBE (red circle) and MCP no Loop TRIBE (background TRIBE, blue circle). (D) Dot plot of data from (C). Individual genes from MCP TRIBE in cells with β-actin-MBS plotted on x axis (red circle), MCP TRIBE in cells without β-actin-MBS plotted on y axis (blue circle). The RNAs found in both experiments are plotted on the diagonal (purple circle).

### Off-target characterization for CLIP and TRIBE

To characterize these targets further, we performed the CLIP and TRIBE without MCP or without the stem loops. CLIP of cells without MCP-GFP generated nearly 200 significant peaks, reproducing 40% of the peaks present in cells with MCP-GFP (Fig. 2B). These peaks most likely reflect nonspecific antibody interactions. In support of this interpretation, there was a five-fold increase in transcript targets when polyclonal antibodies rather than monoclonal antibodies were used (Biswas et al, manuscript in preparation).

In contrast, MCP TRIBE in cells without the β-actin-MBS reproduced only 13% of the with MBS-peaks (Fig. 2C), and there were no transcripts with editing levels comparable to the 27 β-actin-MBS sites, e.g., more than 4 editing sites (Fig. 2D). Interestingly TRIBE showed far fewer off-target peaks compared to CLIP; only 28 transcripts in the entire transcriptome had at least two editing sites.

Since the MCP β-actin-MBS TRIBE target transcripts were highly reproducible, did not have a MCP consensus sequence, were not found in MCP CLIP and were dependent on the presence of the MBS array (they did not appear in MCP-ADAR expressing cells without MS2 stem loops), we considered explanations for these particular off target transcripts.

### Origin of MCP TRIBE off-target signals can be correlated with the genetic locus

One possibility for these reproducible transcripts detected by TRIBE was that they could be proximal to the β-actin-MBS bound ADAR. For instance, if the mRNAs were packaged with β-actin mRNAs in a granule, the mRNAs might come close enough together to facilitate transediting. If this were the case, we would expect the transcripts to share a common feature, such as a zipcode consensus sequence or a function such as motility. However, interrogation of the targets did not turn up a common consensus sequence, typical of the β-actin mRNA zipcode (Patel et al., 2012). Additionally, gene ontology analysis failed to find functions related to β-actin’s role in the cytoplasm (cell motility, migration). We then considered that the targets could be close enough together to undergo trans-editing in the nucleus; perhaps in some nuclear domain. To this end, he off-target transcripts edited by MCP TRIBE were interrogated to determine if β-actin-MBS mRNA could be in proximity within the nucleus. (Fig. 3A).

**Fig. 3.**
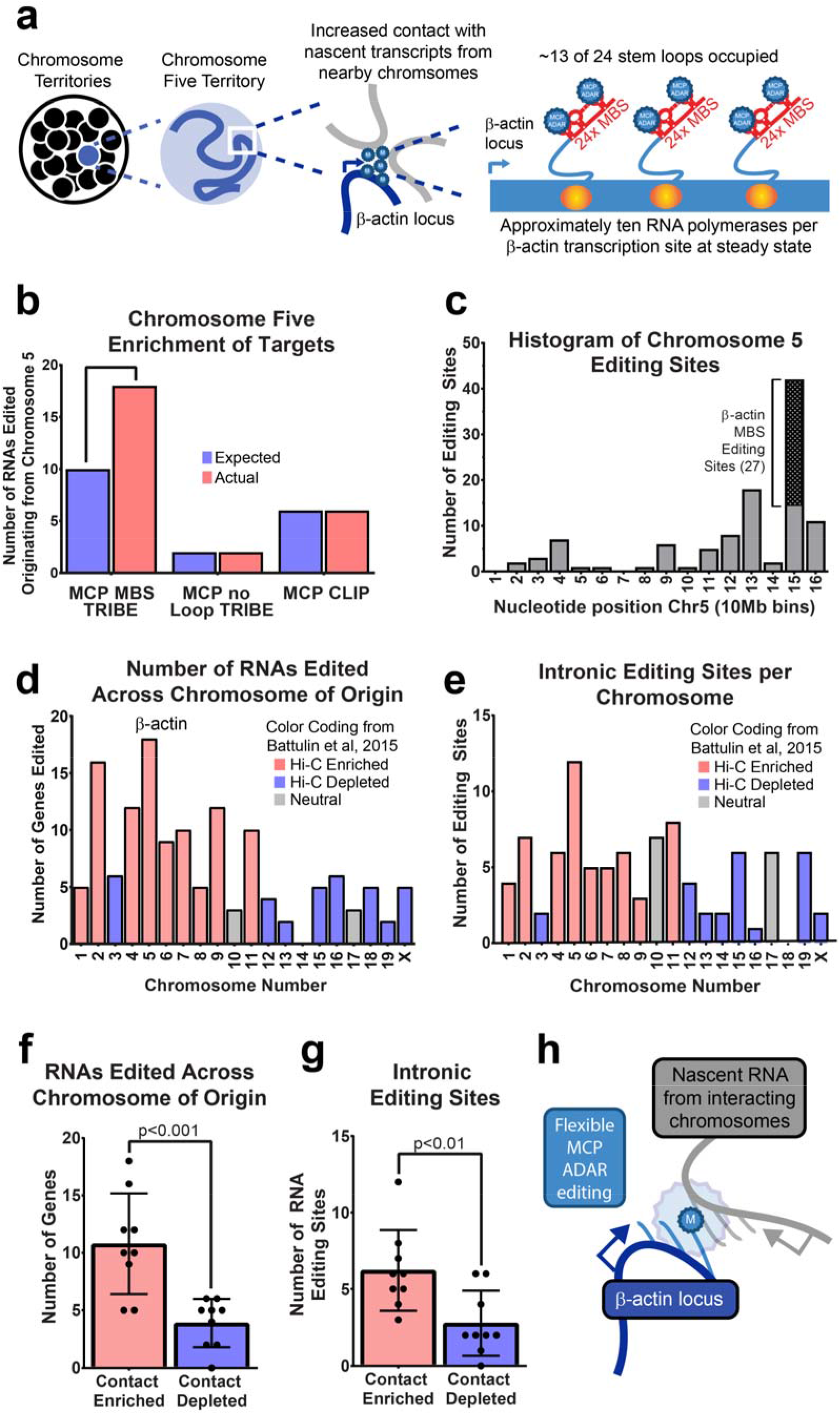
Increased presence of editing sites on chromosome 5 and increased density of editing around the β-actin locus. (A) Schematic of cell nucleus (large circle) containing individual chromosomes within their territories (smaller black circles with chromosome five highlighted in blue). Enlarged view of chromosome five territory showing a snapshot of DNA (blue line), as well as region of flexible movement (blue circle). Enlarged view of ActB gene locus, showing transcription of β-actin mRNA and loading of MCP ADAR near the site of transcription and chromatin contacts. Schematic showing increased loading of MCP ADAR onto actively transcribing RNA. (B) Bar graph showing enrichment (red) vs expected (blue) number of sites for chromosome 5 across experiments. (C) Bar graph showing number of editing sites (y axis) along chromosome 5 (x axis, 10Mb bins). Editing at the bin containing the β-actin-MBS is represented as a hashed box. (D) Bar graph showing number of RNAs edited organized by chromosome of origin. Chromosomes are colored by Hi-C contacts with Chromosome 5 (Data from(Battulin et al., 2015)). Chromosomes enriched for contacts are colored in red, chromosomes depleted in contacts are colored in blue. (E) Number of intronic editing sites organized by chromosome of origin. Chromosomes are colored by Hi-C contacts with Chromosome 5 (Data from (Battulin et al., 2015)). Chromosomes enriched for contacts are colored in red, chromosomes depleted in contacts are colored in blue. (F) Data in D represented as a bar graph. Chromosomes were group by their relationship to Chromosome Five in MEFs, either into contact enriched (red) or contact (depleted). Unpaired, two tailed t-test was used to determine significance. (G) Data in E represented as a bar graph. Chromosomes were grouped by their relationship to Chromosome Five in MEFs, either into contact enriched (red) or contact (depleted). Unpaired, two tailed t-test was used to determine significance. (H) Schematic of proposed model for chromosome contacts. Transcription sites (TS, circles) often loop away from chromosome of origin and intermingle with other active loci. The β-actin-MBS (blue stars) recruits MCP-ADAR at time of transcription and marks nearby active loci.

Surprisingly, many of these transcripts were on the same chromosome as the β-actin-MBS. When normalizing for size, chromosome 5 genes were edited more often than expected in MCP TRIBE data when the β-actin-MBS was present. In contrast, there was no enrichment when the experiment was repeated without the MBS array or when direct protein RNA contacts were queried in MCP CLIP (Fig. 3B).

Spatial interactions would be expected to affect regions proximal to the β-actin locus more than distal parts of chromosome 5. Indeed, higher numbers of chromosome editing sites were near the β-actin-MBS array (Fig. 3C). After addressing cis-contacts on chromosome 5, transcripts from other chromosomes were queried to determine if trans-interactions with chromosome 5 could explain some of the other identified targets. Previously published Hi-C inter-chromosomal contacts from MEFs found chromosome 5 to have enriched contacts with chromosomes, 1,2,4,5-9,11 and depleted for contacts with chromosomes 3,12-16,18,19,X (Battulin et al., 2015). The most enriched chromosome contacts in this dataset were 2, 7 and 11 with chromosome 3 being notable as an example of a large chromosome with depleted contacts. Consistent with these Hi-C data, edited RNAs originated more frequently from chromosomes predicted to have more chromosome 5 contacts (Fig. 3D, 3F, Sup Fig. 2A, Sup Fig. 2B, Sup Fig. 2C). and less frequently from chromosome 3. The latter also indicated that chromosome size was not a major confounding factor.

As intronic reads even from in poly A selected RNA-seq were recently shown to result from unspliced nascent precursor mRNAs, editing of nearby nascent pre-mRNAs should be enriched in intronic reads (Manno et al., 2018). Indeed, chromosomes with enriched chromosome 5 contacts had more intronic editing sites (Fig. 3E, 3G). Further validation was performed by interrogating whether the genes identified as being close to the β-actin-MBS locus had their transcription sites in spatial proximity as has been provided by the intron seq-FISH data (Shah et al., 2018). Of the RNAs edited by MCP TRIBE, 85 of 86 transcription sites interacted with chromosome five as determined by the previously reported seq-FISH results (closer than 500nm). Strikingly, 56 of the 100 genes closest to the β-actin locus on chromosome 5 were MCP TRIBE targets (Sup Fig. 2D). This significant overlap indicates that edited RNAs in the MCP TRIBE experiment represent intra- and inter-chromosomal contacts during transcription.

It became apparent that editing could be a proximity labeling approach that can define a nuclear domain where transcripts spatially interact. By accessing Hi-C data we determined that transcripts that were edited on other chromosomes were likely to be nearby (Fig. 3H, Sup Fig. 2A). In contrast to chromosome conformation capture, where high sequencing depth is required to achieve megabase resolution, RNA editing is able to measure inter-chromosomal interactions with individual transcript resolution (Sup Fig. 2A).

## Discussion

By adapting TRIBE to mammalian cells and converting it to a single plasmid system, a number of advantages were evident. Mammalian TRIBE is an antibody independent approach that can be performed on 1000x fewer cells (samples were generated from fewer than 10k cells) than CLIP (10-20 million cells required) or chromosome conformation capture based approaches (requiring >10 million cells). The sequencing depth required for TRIBE is similar to either CLIP or chromatin conformation capture variants, but TRIBE library preparation is upon total RNA. By retaining the sequence complexity of the cellular transcriptome, fewer PCR cycles are required for library amplification thus leading to increased sequencing efficiency. Significantly more sequenced TRIBE reads are usable than the most efficient CLIP approaches (Sup Fig 3A, 3B).

During the process of gel excision, CLIP tags are limited in size due to the additional mass of the protein-RNA complex, 100nt of RNA adds approximately or 32kDa of mass to the protein-RNA smear. Post processing, most CLIP tags are ~50nt, in contrast, TRIBE can utilize longer RNA sequencing reads (150nt here) which increases its ability to uniquely map reads (Sup Fig 3A). This allows TRIBE to discover the entire repitore of RNA targets, as evidenced by saturated target discovery and RNA editing (Sup Fig 3B, 3C). Additionally, the molecular biology required for TRIBE is radiation free and limited to cell sorting, RNA isolation and standard RNA library preparation, processes that are routine in many labs and can be completed in as few as three days.

### Definition of background

Until recently, limited alternative approaches to CLIP existed. One prior approach was pertinent to the study of splicing factors. In this context, CLIP was assisted by correlating binding sites with sites of alternative splicing, as determined by RNA seq. Ultimately, multiple transcriptome wide approaches should be integrated to evaluate the targets of RNA binding proteins (Rahman et al., 2018).

Prior work comparing CLIP with RNA tagging in yeast found ~50% overlap between the two approaches (Lapointe et al., 2015, 2018). Similar percentages were found when comparing CLIP and TRIBE in *Drosophila* (McMahon et al., 2016). However, most comparisons have been performed on RNA binding proteins that bind to a significant percentage of cell transcripts (often > 50%), thus confounding the value of percentage overlap. The large number of bound targets also makes it challenging to identify the origin of these differences. By using the MCP MBS system we were able to perform CLIP and TRIBE with only one true target within the cell. The small number of discovered targets in each technique (<200 of 15k transcripts) avoided spurious overlap and readily identified overlap as well as false positives.

For CLIP, the use of KO controls reveals nonspecific RNA binding to antibodies. A similar issue has been described in ChIP-seq data from both *Drosophila* and yeast, leading to the presence of “phantom peaks” that are present even when the antigen is not (Jain et al., 2015; Teytelman et al., 2013).

### Identification of chromatin contacts and transcription domains

The small number of MCP TRIBE targets and higher signal to noise allowed us to discover the cause of off-target transcripts. When multiple copies of MCP-ADAR were tethered to a highly transcribed 24xMS2 locus, editing also affected nearby transcripts, both on its own chromosome and nearby chromosomes. This is likely assisted by the frequent transcription of the β-actin mRNA in MEFs. At steady state >80% of the cells have more than one active β-actin transcription site and each site averages 10 ±3 molecules of β-actin mRNA at a given moment (Kalo et al., 2015). As 13 molecules of MCP are on average loaded onto each molecule of β-actin mRNA(Wu et al., 2012), the accumulation of ~130 molecules of MCP-ADAR at the β-actin transcription site (chromosome 5) likely enhances the amount of RNA editing of nearby transcripts. Additionally, β-actin borders a super enhancer in both mouse and human cells (Khan and Zhang, 2016), likely furthering its interactions with other genes and chromosomes.

This would only occur if the transcribing loci from nearby chromosomes shared intimate contacts with the β-actin transcription site. Inter-chromosomal contacts have been identified by Hi-C (Battulin et al., 2015; Lieberman-Aiden et al., 2009) and further supported by transcriptome wide RNA FISH studies in MEFs that observed nascent transcripts looping away from the chromosomal DNA (on average 0.8 ± 1.1μm away). This allowed intermingling with other nascent transcripts even from multiple chromosomes (Shah et al., 2018). Due to the limited resolution of ligation based approaches (Hi-C) and optical microscopy (intron seq-FISH), specific interactions at the nucleotide level could not be previously resolved. Additionally, the requirement for edited RNA focuses on loci that are simultaneously transcribing and should reduce contributions from many other sources of DNA/DNA interactions. This functional filter may simplify the extensive interchromosomal contacts that have been observed with other approaches.

Recently, proteins that interact with the β-actin-MBS were profiled using MCP fused to the biotin ligase BirA. Interestingly, several nuclear proteins and chromatin components were discovered even after control subtraction, suggesting a similar co-transcriptional contribution to RNA-protein profiling when the MBS array is used (Mukherjee et al., 2019). However, TRIBE differs in the mechanism of transcriptional marking: the enzyme is affixed to the transcript rather than an interaction that occurs by chance diffusion of a reactive intermediate. Therefore, the transcripts must be in closer physical contact rather than simply nearby.

### Physiological considerations

The fact that nascent transcripts intermingle makes it possible to share factors, for instance the effective concentration of splicing factors would be increased by the proximity of the transcripts undergoing splicing. This should significantly increase reaction rates, an explanation for why *in vivo* splicing is so efficient. ADAR may possibly edit excised introns but since these are shortlived compared to the exonic sequences, it is more likely that the editing occurs before or during splicing. Effects on splicing rates may be testable by assaying a splicing reporter placed near highly transcribed and rapidly spliced genes vs further away from other spliced genes.

The distance from the MCP (PDB: 2BU1) bound to the stem loops to the editing pocket of the ADAR (PDB: 5ED1) is only a few nanometers (<10nm, similar to the length of 30 nucleotides of folded RNA), indicating that transcripts come very close to one another. This may be due to the looping of nascent chains into a constrained volume, or nascent chains may explore a much larger volume so they eventually come into contact with each other. In either case, collisions of these RNA strands do not result in entanglements, and examples of trans-splicing are rare. This could best be accomplished if the nascent chains assume a minimal spherical volume rather than a linear dimension, so editing would then occur where these surface regions of exposed nucleotides come into contact.

Because the ADAR is an enzyme, it requires some time to find and modify the appropriate adenosine. This has been estimated to be about 24 seconds (Kuttan and Bass, 2012). Therefore, ADAR must stay in contact with its substrate longer than would be required for CLIP, where crosslinking is instantaneous. This would provide a filter for more persistent interactions and possibly more physiologically relevant ones (McMahon et al., 2016).

The discovery of chromatin contacts by MCP TRIBE suggests a number of future directions. Large nuclear foci consisting of nascent transcripts have been previously observed (Fay et al., 1997) and more recent studies have proposed a transcriptional hub model where multiple transcribing loci spatially organize (Cho et al., 2018; Hnisz et al., 2017). Several theories have been proposed for the function of transcriptional hubs, in particular our data support the role of transcription factors in hub formation (Liu and Tjian, 2018). Specifically, half of our MCP TRIBE targets contain upstream binding sites for serum response factor (Roider et al., 2009), further work will determine if additional parameters such as processing and splicing rate are correlated across contacts and if this coordination is mediated by specific transcription factors.

Transcription hubs change during stem cell differentiation and have been shown to contain multiple copies of both mediator and RNA polymerase II (Cho et al., 2018). To date, it is still not known which genes are actively transcribing within these transcriptional hubs. The highly multivalent nature of proteins within the hub, e.g., Mediator, make it an attractive target for future studies of chromatin contacts with TRIBE. Additionally, the discovery of enhancer RNAs makes them an attractive target for MS2 tagging and subsequent TRIBE to determine a list of functional RNA targets. The development of MS2 tagged gene libraries and CRISPR mediated integration will further allow chromatin contacts to be studied by MCP TRIBE.

## Materials and Methods

### Modifications from previously published TRIBE protocol

When adapting the technique from Drosophila (McMahon et al., 2016; Rahman et al., 2018) a number of changes were made. The first being that the two plasmid system required for TRIBE (one plasmid carrying the RBP fused to ADAR, the other carrying GFP as a marker) was combined onto a single multicistronic vector by using the p2A system. The equimolar expression for the upstream RBP-ADAR and downstream GFP cistrons, allowed for accurate quantification RBP expression levels during FACS sorting (Lo et al., 2015). Additionally, competition from the endogenous RBP was removed by performing all experiments in KO backgrounds. These same lines were also used as background controls for CLIP studies, highlighting antibody specific contributions to downstream CLIP analysis. To further increase the signal to noise of RNA editing, two dADAR point mutations were used. The hyperediting mutant of dADAR (E488Q) had previously been shown to be 10x more active (Kuttan and Bass, 2012) and had less sequence bias (Xu et al., 2018). To further increase signal, S458 was synonymized to stop auto inactivation of RNA editing (Palladino et al., 2000).

### Plasmid construction

tdMCP-stdGFP (Addgene # 98916) was modified to generated the vectors for this study. The first GFP was excised and replaced with dADAR E488Q with synonymized serine 458 codon (synthesized by IDT). A p2A sequence was added to the end of dADAR by PCR. Once this vector was made, MCP was replaced by either mCherry or ZBP1.

### Cell culture and transfections

Primary MEFs were isolated from the E14 MBS embryos or ZBP1 KO MBS embryos(Katz et al., 2012). Primary MEFs were then immortalized by transient transfection with a plasmid expressing SV40 large T antigen (Addgene #21826). Single cell clones were then isolated by limiting dilution. MEFs were continuously cultured in 10% DMEM (4.5g/L, Corning) supplemented with penn/strep (Gibco) and 10% FBS (Atlanta Biologics). Transient transfection was performed with JetPrime (Polyplus) as per manufacturer instructions 12 hours before sorting.

### FACS sorting

Cells were trypsinized and suspended in sorting buffer (DPBS, 1% BSA supplemented with DAPI) and passed through a single cell mesh filter. FACS was performed by selecting for GFP positive DAPI negative single cells on a BD Aria II instrument. At least 10k GFP positive cells were directly FACS sorted into 800uL Trizol (Invitrogen) and RNA isolation was performed as per manufacturer’s instructions.

### RNA isolation and library prep from MEFs

GFP+ DAPI-MEFs were FACS sorted directly into Trizol and RNA was extracted, quantified with qubit HS RNA assay and integrity was visualized with a RNA pico chip (Agilent Bioanalyzer). Samples with no apparent degradation were used to prepare stranded libraries (NEB Ultra II Stranded RNA seq library prep kit with poly A RNA isolation module) with 150ng RNA input per sample (as quantified by qubit RNA HS assay). Final libraries were amplified with 1-2 rounds of PCR less than the manufacturers recommendation and checked on the bioanalyzer for appropriate size distribution. Library concentrations were then quantified by qubit dsDNA HS assay, qPCR and Agilent bioanalyzer (the latter two of which were performed by Novogene and used for multiplex library normalization prior to sequencing).

### Sequencing and analysis of TRIBE data

Samples were then combined and multiplexed across multiple lanes for HiSeq 4000 runs (performed by Novogene). Paired end sequencing was performed with 150bp reads at a depth of at least 27.5Gb per sample (this corresponded to at least 40 million sequenced read pairs per sample). The data was analyzed following the publicly available computational pipeline [https://github.com/rosbashlab/HyperTRIBE/] developed here(Rahman et al., 2018) with a few modifications described in this section. The github repository has been updated to incorporate the modifications. Reads from sequencing libraries are trimmed and aligned to the transcriptome, which was modified to include the additional MS2 sequence at the β-actin locus as a separate contig. Subsequently, the nucleotide frequency at each position in the transcriptome is recorded from aligned reads. For each nucleotide in the transcriptome, the TRIBE RNA nucleotide frequency is compared with the wild type mRNA library (wtRNA) nucleotide frequency to identify RNA editing sites. For a legitimate edit site, the frequency of A is greater than 80% and the frequency of G is 0.5% in the wtRNA and the frequency of G is greater than 0 in the TRIBE RNA (using the reverse complement if annotated gene is in the reverse strand). There are two modifications compared to the previous version of the TRIBE pipeline. Modifying the nucleotide frequency of G in wtRNA from 0 to 0.5%allows very low level sequence heterogeneity in wtRNA and does not disrupt the identification of legitimate editing sites in deeply sequenced libraries. We used an editing threshold of at least 5% in each replicate, as lowering the threshold from 10% did not significantly alter the percentage of TRIBE target genes with consensus sequence overlap. The TRIBE edit sites are required to be present in both replicates and any sites that overlap with catalytic Adar alone edit sites with at least 1% editing are removed. Finally, a list of transcripts is created by tabulating all transcript that contain edit sites.

### CLIP protocol (From (Cho et al., 2012)) with the following adaptations

15cm dishes of MEFs were washed with ice cold DPBS, after aspiration and addition of 12mL ice cold DPBS, cells were irradiated with 300mJ/cm of 254nm UV light (BioRad GS Gene Linker). After irridation, each 15cm dish was scraped and collected into a 15mL falcon tube. Cells were pelleted at 500g for 5 minutes and then the pellets were stored at −80 until use.

For Immunoprecipitations, a mix of monoclonal Anti-GFP antibodies from Sigma Aldrich / Roche (11814460001) were used with protein G dynabeads. All washes were performed using a magnetic stand.

### Sequencing and analysis of CLIP data

CLIP libraries were sequenced with 75bp SE reads on NextSeq500 using the high output mode. Both replicates were concatenated before peak calling. Both traditional peak calling with Clip Tool Kit (CTK) (Shah et al., 2017) [https://zhanglab.c2b2.columbia.edu/index.php/Standard/BrdU-CLIP_data_analysis_using_CTK] as well as CIMS analysis (Moore et al., 2014) were performed.

### Interchromosomal Hi-C Analysis

The comparison to MEF Hi-C data to determine interchromosomal enrichment / depletion done using two different data sets. Chromosome five contacts with all other chromosomes were considered and data was binned into three groups (enriched, depleted or neutral).

Both data sets were mapped using the iterative correction outlined(Imakaev et al., 2012) and gave consistent results. The first data set utilized previously published data(Battulin et al., 2015) and was analyzed by hiclib software. A separate MEF Hi-C experiment was analyzed using the Juicer suite of tools. Interchromosomal contacts were isolated from a .hic file containing high quality mappings (>30).

The expected number of interactions between each chromosome pair (chromosomes i and j for example) was calculated(Lieberman-Aiden et al., 2009). Reads originating from between two different chromosomes (interchromosomal) were considered.

The fraction of interchromosomal reads coming from chromosome i* fraction of interchromosomal reads coming from chromosome j* total number of interchromosomal reads

The expected number of interactions represents chances of a read randomly occurring between chromosome i AND chromosome j based on the total number of interchromosmal reads generated.

Enrichment was then calculated by taking the number of interactions observed between chromosomes i and j and dividing by the expected value. Chromosomes enriched above the average for chromosome five were considered enriched and those below average were considered depleted.

### Data availability statement

All raw sequencing data and identified RNA editing sites are available for download from NCBI Gene Expression Omnibus (http://www.ncbi.nlm.nih.gov/geo/), accession number ####. Plasmids encoding mCherry-ADAR (Addgene number ####) and MCP-ADAR (Addgene number ####) are available at Addgene. All other relevant data are available from the authors upon request.

## Acknowledgements

The authors would also like to thank Charles Query, members of the Singer Lab, and the Rosbash Lab for their helpful discussions and comments on the manuscript. RHS was supported by R01NS083085 and U01DA047729. MR was supported by the Howard Hughes Medical Institute and R01DA037721. JB was supported with funding from an MSTP Training Grant T32GM007288 and predoctoral fellowship F30CA214009. FACS sorting was performed with core support from NIH P30CA013330.

## Author Contributions

JB designed, performed all experiments and generated all data used in this study, RR and VG processed the RNA seq data and was assisted by JB who performed further analysis of the processed data. JB and RHS drafted the manuscript, RR, RHS and MR edited the manuscript. RHS and MR supervised the research.

