## Supplementary Material for "MS2-TRIBE evaluates protein-RNA interactions and nuclear organization of transcription by RNA editing"

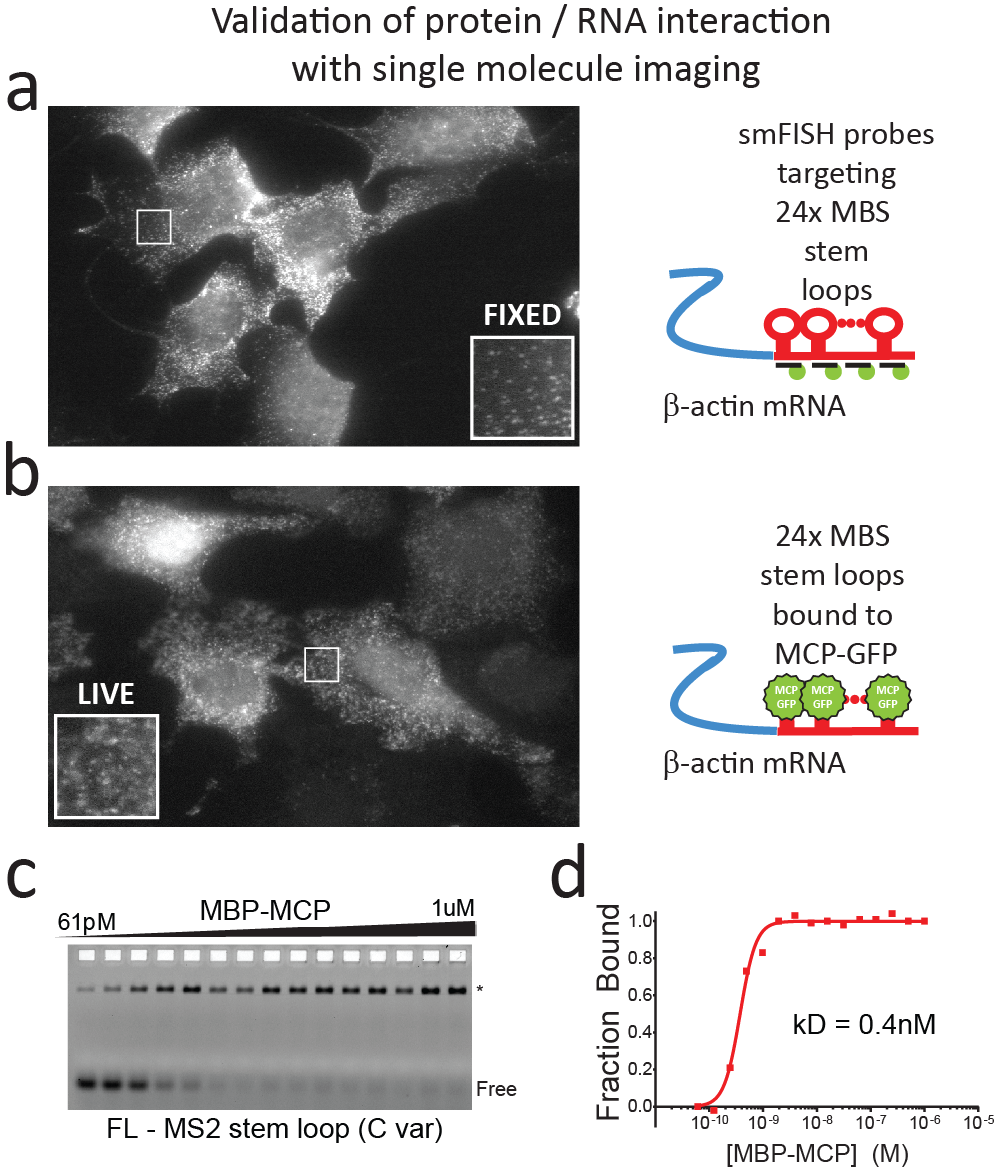

**Supplementary Fig 1. Single molecule imaging and biochemical characterization of the MCP MBS complex**

(a) Immortalized MEFs from the β-actin-MBS mouse were fixed and smFISH was performed using probes against the stem loop array. Single molecules of β-actin mRNA were visualized as bright, uniform intensity spots in the cytoplasm (insert).
(b) Live imaging of the immortalized β-actin MBS MEFs was performed by stably infecting the cells with MCP-GFP and FACS sorting the GFP positive populations, single molecules were again visualized and had uniform fluorescence intensity (insert).
(c) Representative electrophoretic mobility shift assay (EMSA) MS2 stem loop RNA. The filled triangle represents a 1:1 serial dilution of MCP. The RBPRNA complex (*) and free RNA (FREE) are labeled.
(d) Quantification and fit to the Hill equation of EMSA results for MCP (red solid line).

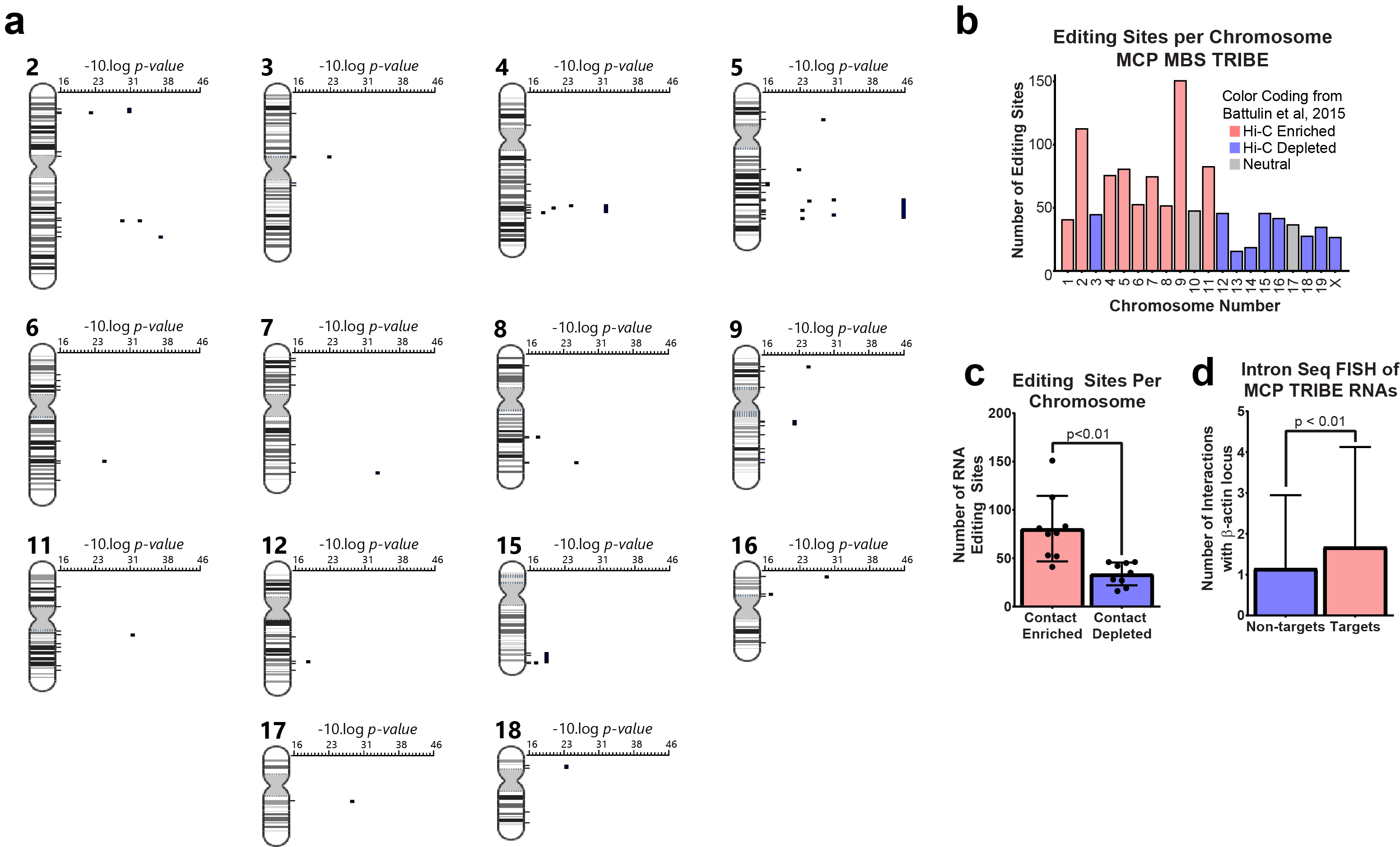

**Supplementary Fig 2. MCP TRIBE shows significantly edited regions of DNA corresponding to chromosome contacts**

(a) Positional gene enrichment of MCP TRIBE results in β-actin-MBS cells. To left are chromosome numbers with diagrams beneath. Significantly enriched regions are displayed as black boxes from left to right, in order of increasing significance (-10 log pvalue).
(b) Bar graph showing number of editing sites organized by chromosome of origin. Chromosomes are colored by Hi-C contacts with Chromosome 5 (Data from (Battulin et al., 2015)). Chromosomes enriched for contacts are colored in red, chromosomes depleted in contacts are colored in blue.

**Actb 24x MS2 Stem Loop**

(c) Data in B represented as a bar graph. Chromosomes were grouped by their relationship to Chromosome Five in MEFs, either into contact enriched (red) or contact (depleted). Unpaired, two tailed t-test was used to determine significance.
(d) Bar graph showing average number of contacts (as calculated by seq-FISH interaction matrix) for MCP-TRIBE targets vs non targets with the region surrounding β-actin-MBS.

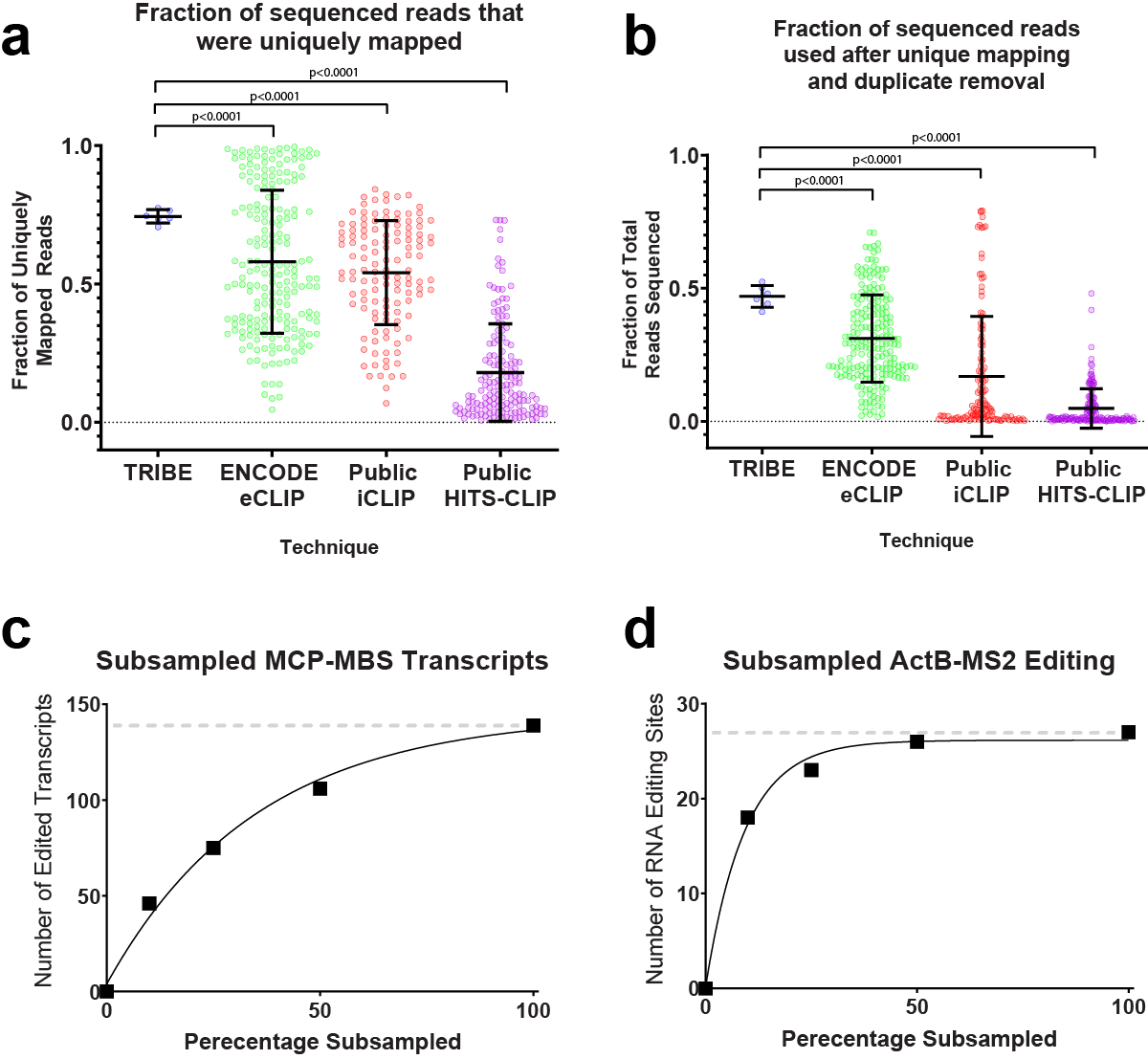

**Supplementary Figure 3. TRIBE libraries are more complex and more efficiently sequenced than CLIP libraries, allowing discovery of targets to be saturated**

(a) Percentage of reads in FASTQ file that were uniquely mapped to the genome. Individual points represent individual experiments or biological replicates. P values calculated using two-tailed Welch’s t-test. Processed CLIP data from (Van Nostrand et al., 2016).
(b) Percentage of reads in FASTQ file that are retained after complete processing. Major RNA seq processing steps include unique mapping and PCR duplicate removal. Individual points represent individual experiments or biological replicates. P values calculated using two-tailed Welch’s t-test. Processed CLIP data from (Van Nostrand et al., 2016).
(c) Scatter plot showing number of edited transcripts as a function of random subsampling from the aligned reads. X axis, subsampled percentages (100%, 50%, 25%, 10%, 0%). Y axis, number of RNAs in the transcriptome that were edited by MCP-TRIBE (black boxes). Solid black line represents nonlinear fit of the data (one phase association).
(d) Scatter plot showing number of RNA editing sites on β-actin-MBS transcript a function of random subsampling from the aligned reads. X axis, subsampled percentages (100%, 50%, 25%, 10%, 0%). Y axis, number of RNAs in the transcriptome that were edited by MCP-TRIBE (black boxes). Solid black line represents nonlinear fit of the data (one phase association).
